# Multi-SpinX: An Advanced Framework for Automated Tracking of Mitotic Spindles and Kinetochores in Multicellular Environments

**DOI:** 10.1101/2024.04.03.587736

**Authors:** Binghao Chai, Christoforos Efstathiou, Muntaqa S. Choudhury, Kinue Kuniyasu, Saakshi Sanjay Jain, Alexia-Cristina Maharea, Kozo Tanaka, Viji M. Draviam

## Abstract

SpinX, an AI-guided spindle tracking software, allows the 3-dimensional (3D) tracking of metaphase spindle movements in mammalian cells. Using over 900 images of dividing cells, we create the Multi-SpinX framework to significantly expand SpinX’s applications: a) to track spindles and cell cortex in multicellular environments, b) to combine two object tracking (spindle with kinetochores marked by centromeric probes) and c) to extend spindle tracking beyond metaphase to prometaphase and anaphase stages where spindle morphology is different. We have used a human-in-the-loop approach to assess our optimisation steps, to manually identify challenges and to build a robust computational pipeline for segmenting kinetochore pairs and spindles. Spindles of both H1299 and RPE1 cells have been assessed and validated for use through Multi-SpinX, and we expect the tool to be versatile in enabling quantitative studies of mitotic subcellular dynamics.

## 1 Introduction

The advancement of deep learning techniques has spurred a rise in the utilisation of computational image analysis in cell biology [1, 2] and drug discovery studies [3]. While computational methods have dramatically transformed the automated analysis of static microscopy images [1, 2, 4, 5], their application to time-lapse microscopy videos for studying the structural dynamics of objects in 3D space and through time is still emerging [6–9]. In addition to computational advances, the rapid development in live-cell imaging technologies, enabling the capture of dynamic subcellular processes with remarkable clarity, has resulted in the generation of large-scale, high-resolution and time-lapse movies, presenting both an opportunity and a challenge for automated high-throughput image analysis [2, 10]. Although an increasing amount of imaging data provides an abundance of detailed sub-cellular information, their manual analysis is not only a tedious task, prone to biases, but also an underutilisation of the invaluable skills of experienced researchers. The development of new computational architectures tailored to time-lapse movies of specific subcellular structures holds the promise for significantly advancing the quantitative analyses of subcellular and cellular dynamics. However, automated analysis of live-cell movies remains a challenging problem in image analysis due to discontinuities in time-lapse movies [2, 10].

Imaging across scales of space and time is a well-recognised challenge in high-resolution fluorescence microscopy arising due to phototoxicity that dividing cells can not withstand for long periods of time [11–13]. Dividing mitotic cells with condensed DNA are extremely sensitive to light, and this disallows continuous imaging, making it challenging to track dynamically moving objects in mitotic cells.

During mitosis, the microtubules of the mitotic spindle are tethered to chromosomes by a specialised macromolecular structure, the kinetochore that assembles on the centromeric region of chromosomes [14]. The mitotic spindle is constantly changing shape and it undergoes movements independent from the kinetochores that are also moving within the spindle. The tracking challenge becomes computationally intensive due to either the crowding of objects within cells or the high-throughput nature of the study presenting many dividing cells next to each other. Managing these challenges of tracking multiple spindles in an image or multiple kinetochores (smaller objects) within a spindle requires breaking down the complexity into smaller individual units. This would entail isolating and tracking the larger objects (spindles) first and then focusing on tracking the smaller objects (kinetochores) within them.

Here we present a framework to track multiple spindles and kinetochores, taking advantage of SpinX, an AI-guided segmentation and 3D-tracker of metaphase spindles [7]. Using high-quality image datasets alongside cutting-edge computational methods, SpinX can accurately track 3D movements of individual mitotic spindles during metaphase [7]. SpinX was limited to analysing single cells and spindles in metaphase. In response to the growing demand for a more adaptable tool, suitable for high-throughput studies, we have now developed a framework to enhance the capabilities of SpinX. The iteration we introduce in this paper, termed Multi-SpinX, represents a significant expansion of our original framework. It is specifically engineered to broaden SpinX’s functionality to include not just single-cells, but also multicellular and multi-content imaging scenarios. The primary objective of Multi-SpinX is to broaden the usability of SpinX, making it applicable to a wider variety of datasets beyond metaphase spindles. This expansion includes refined mathematical modelling, enabling the precise, automated tracking of multiple mitotic spindles across diverse cellular environments. Using experienced and naive users, we evaluate the extent to which our expanded new solution is robust, and we optimise it for analysing complex live-cell movies. In summary, the expanded Multi-SpinX optimised for high-throughput studies is expected to save invaluable expert researcher time and pave the way for new explorations into the dynamic movements of chromosomes, kinetochores and mitotic spindle with an unprecedented level of detail.

## 2 Results

### 2.1 Computational framework to track movements of multiple spindles and kinetochores in a crowded environment

To adapt the existing spindle tracker, SpinX [7], for high-throughput (multiple spindles) and multi-channel (objects other than spindles) studies, we develop two image processing pipelines that can be integrated with SpinX. The pipeline for detecting multiple spindles, and tracking them individually, uses a Hungarian algorithm with Euclidean distance as a loss function [15] and crops individual objects for 3D image segmentation and modelling using SpinX (Fig 1.A, top row). The second pipeline combines the tracking of spindle movements with additional objects. Here we set out a kinetochore tracker pipeline that detects and tracks sister kinetochore pairs that move and separate during cell division (Fig 1.A, bottom row).

**Fig. 1:**
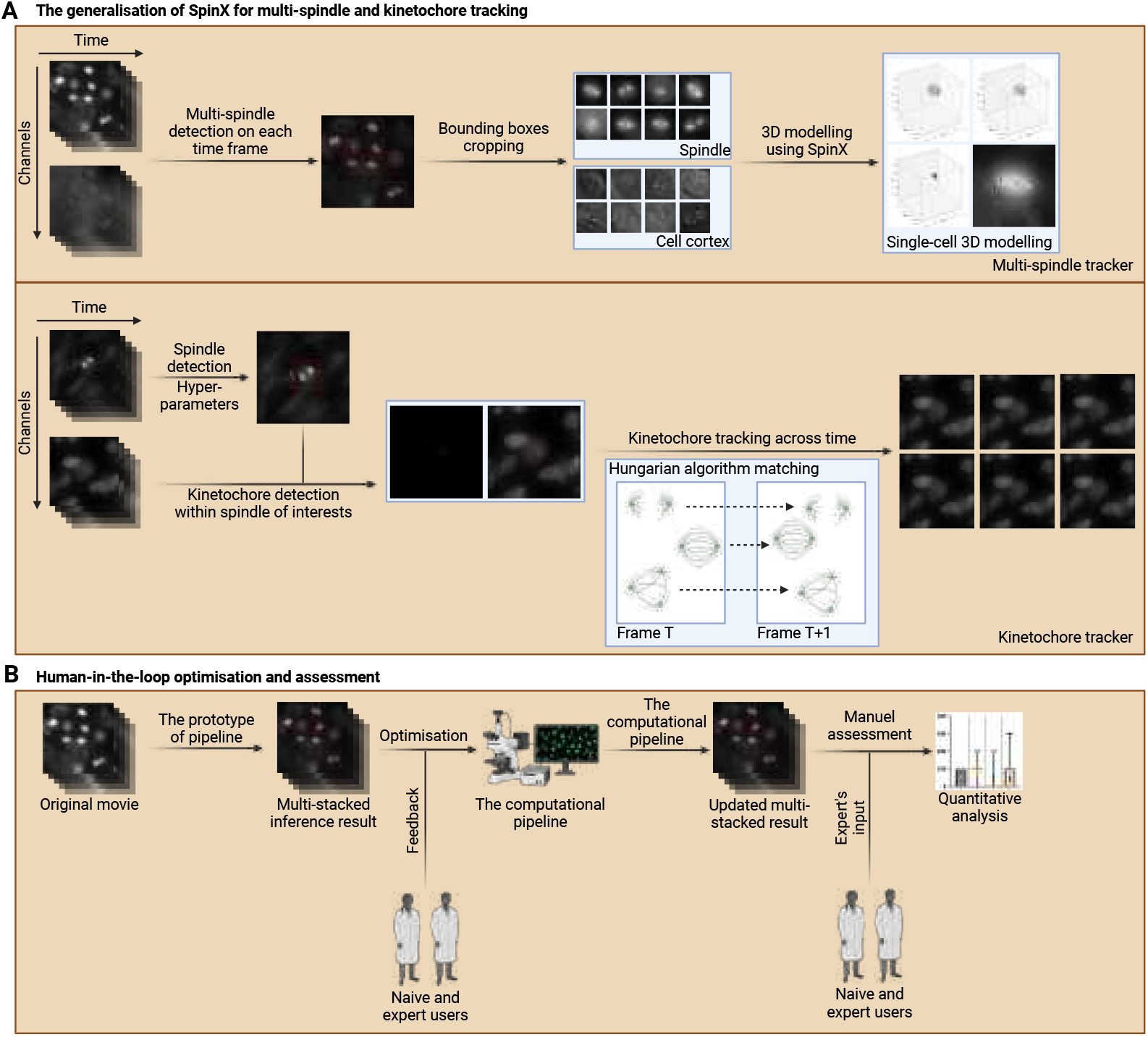
Multi-SpinX framework using two pipelines, a multi-spindle tracker and a kinetochore tracker. (A) Computational pipelines for the multi-spindle and kinetochore tracking. In Stage 1 of both pipelines, Watershed segmentation is applied to generate the bounding box around objects of interest (red boxes). Hungarian algorithm [15] with Euclidean distance as the loss function is then employed for multi-object tracking in Stage 2 of both pipelines. In the multi-spindle tracker, the cropped images of spindles and respective cell cortex are generated before sending them to SpinX for the final step of single-cell 3D modelling. (B) Human-in-the-loop optimisation and assessment. There are two rounds of manual assessments for the computational pipelines. Feedback collated from experts and naive users in the first round of assessment is used to identify computational challenges for pipeline optimisation. As a second and final round of assessment, expert and naive users score tracking accuracy across consecutive time-frames after the optimisation of the two pipelines. This figure was created using BioRender (https://biorender.com/).

The Multi-spindle tracker and Kinetochore tracker were optimised and assessed during their respective development using a human-in-the-loop approach (Fig 1.B) and a large-scale image dataset of multiple spindles and kinetochores within spindles were used for user training and assessment. Together, the Multi-SpinX framework expands SpinX into a multi-spindle tracker and a kinetochore tracker, enabling high-throughput quantifications of spindle and kinetochore movements.

### 2.2 Generalisation of SpinX to track spindles in crowded environments

The AI-guided SpinX software can segment and track metaphase spindle movements, in single cells, visualised using high and low-resolution objectives (60x and 40x magnifications, respectively) [7]. Here, we extend SpinX for high-throughput studies by developing a framework to identify and crop multiple spindles found in a crowded image dataset (Fig 2.A).

**Fig. 2:**
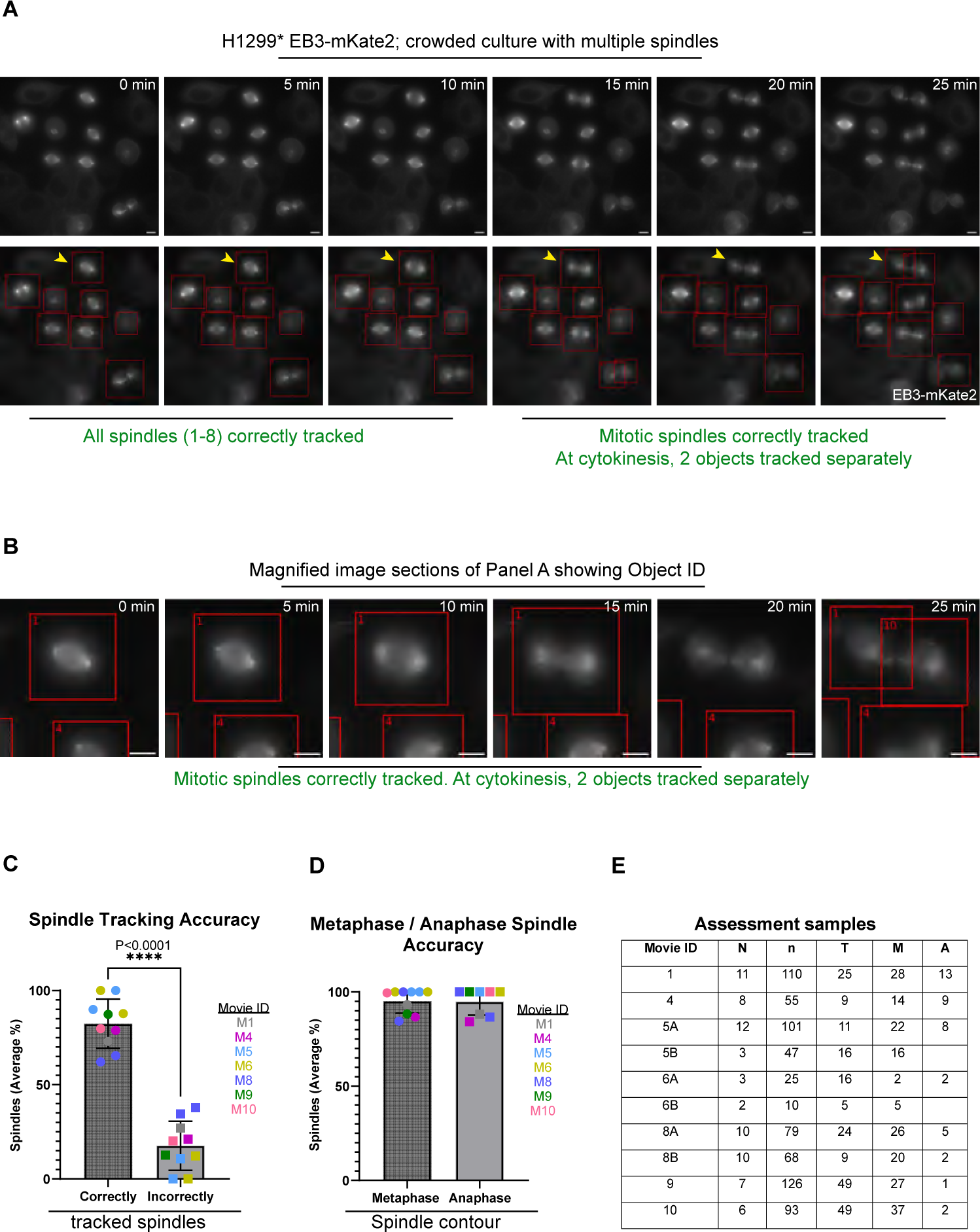
Evaluation of the multi-spindle tracker using time-lapse movies of H1299^*^ EB3-mKate2 cell line. (A) Representative images of multiple spindles in cells expressing EB3-mKate2 (a microtubule-end marker for spindle tracking). Time-lapse images show the dynamic nature of spindle contour, movements and orientation changes over time, captured at specific intervals (0, 5, 10, 15, 20, and 25 minutes). Overlay of tracking results in red bounding boxes are provided with the original frames. This panel serves as a foundational reference for understanding the methodology behind spindle tracking and the visual cues used to assess Multi-SpinX (in green). (B) Magnified images of individual spindle and bounding boxes correspond to areas highlighted with yellow arrows in Panel A showing two bounding boxes at cytokinesis marking the end of cell division (right-most panel). (C) Bar chart of the percentage of spindle bounding boxes correctly tracking spindles across consecutive time frames in 10 movies. Non-mitotic cells were excluded from the analysis. Mean values from 4 separate assessors plotted to quantitatively assess multi-spindle tracker accuracy from prometaphase to telophase and cytokinesis as in panel A. (D) Bar chart assessing the performance of the multi-spindle tracker by comparing its accuracy in tracking metaphase versus anaphase spindles exhibiting different shapes. (E) Table related to sample sizes in panels C and D with Movie ID description showing N (number of different spindles), n (number of events assessed for tracking accuracy), T (time frames indicating movie length) and M/A refers to metaphase anaphase spindles events tracked for accuracy studies. Scale bar: 10 μm.

To identify and crop individual spindles which would make it a suitable input for SpinX, we use watershed segmentation to generate bounding boxes around spindles (Fig 1.A, red boxes) that grow in size as the cell progresses through mitosis through time. Spindles at the very edge of the images are removed from the final dataset to reduce partial tracks. The bounding box could successfully identify cytokinesis by introducing two distinct bounding boxes around the newly formed daughter cells (Fig 2.B).

To assess the accuracy of multi-spindle tracking, correct and incorrectly tracked spindles across consecutive time frames were counted as binary inputs, yes and no, respectively, by a set of four experienced and naive biosciences experts. The first round of assessment yielded poor outcomes specifically in images with less-bright objects (data not shown), and this specific gap was detected by naive and expert users which allowed us to strengthen the computational pipeline by auto-adjustment of movie contrast [16] before the next iteration of tracking assessment.

A total of 213 timeframes (images) with spindles were all assessed by four different assessors and 82.43% accuracy in tracking spindle objects was achieved (Fig 2.C). This is a very high accuracy for this step of the Multi-SpinX framework, aimed at generating SpinX input, since we can crop a continuous stretch of the time-lapse images based on the correct identification of one, two or three consecutive events. In summary, the human-in-the-loop approach allowed rapid optimisation leading to very high accuracy in the framework.

We assessed 72 different spindles evolving their bipolar morphology through time, allowing us to carefully check if the tracking worked well for both metaphase and anaphase spindles (Fig 2.D). Metaphase spindles were present for a longer period in the movie as anticipated while anaphase spindles were disassembled within 5-10 minutes, both are within our expectations [17]. Irrespective of this difference in time and rate of morphology changes, both metaphase and anaphase spindles were tracked with similar accuracies (Fig 2.D and E). These findings indicate that the watershed segmentation-based identification and cropping of single spindles is a good way to study metaphase spindles that progress into anaphase when spindle length elongates with the cell cortex. Based on the manual assessment (Fig 2.A and D)., we conclude the multi-spindle tracker shows high tracking success rates in prometaphase, metaphase and anaphase spindles, highlighting the tool’s capabilities in tracking spindles through multiple phases of cell division.

In summary, the multi-spindle tracker framework can integrate with SpinX to successfully identify spindles in prometaphase, metaphase and anaphase, in crowded high-throughput movies showing multiple dividing cells.

### 2.3 Generalisation of SpinX to kinetochore tracking

Being able to track kinetochores organised within the mitotic spindle can help investigate and understand kinetochore lesions that lead to the incorrect number of chromosomes in prematurely ageing or diseased cells [18]. So, we aimed to combine the spindle tracker with the kinetochore tracker to measure kinetochore movements in relation to spindle movements.

In RPE1 cells treated with SiR-Tubulin dye (a tubulin and spindle marker [19]), we use a CRISPR probe-based tracking of centromeric DNA to label a single chromosome as GFP-labelled pair of particles (leading to labelling two of the kinetochore pairs in each cell). As in Fig 2.A, the spindle tracker is first used to identify large objects to define the bounds of the tracked spindles (Fig 3.A). Within the bounding box (this is also the region of interest for kinetochore detection), the search for centromere/kinetochore particles was constrained which reduced the undesired influence of artefacts or bright objects from neighbouring cells.

**Fig. 3:**
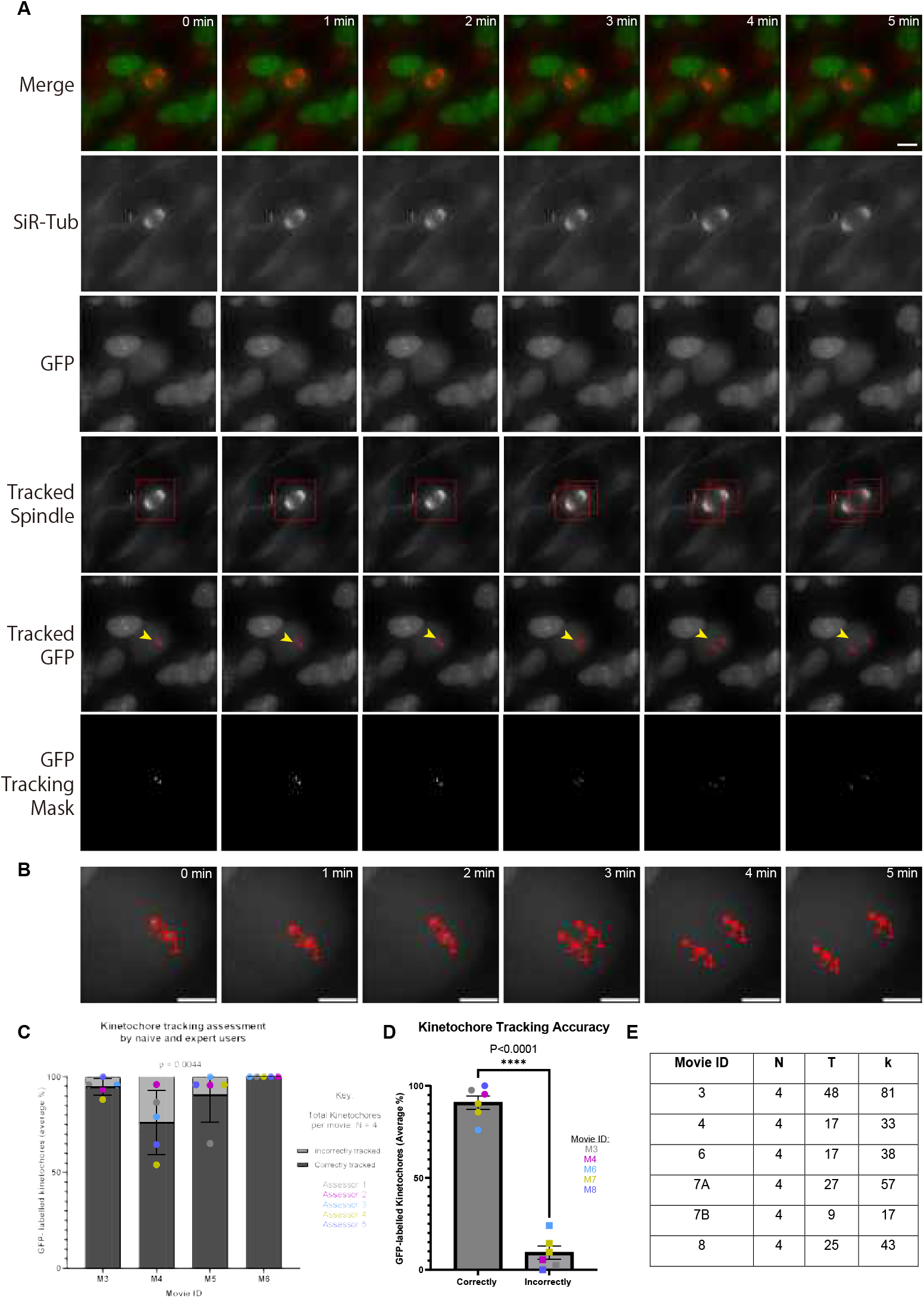
Evaluation of the kinetochore tracker in cells labelled with centromere markers shows the generalisability of the tracker to different chromosomes. (A) Time-lapse images of consecutive frames illustrate the complex process of kinetochore tracking overlaid with spindle tracking. The frames are organised as follows: the original movie as a merge of the spindle (red) and kinetochore marker (centromere marker for chromosome 16, in green) displays the representative consecutive frames (first row), SiR-Tubulin (SiR-Tub) labelled channel highlights the spindle structure (second row), GFP channel highlights the kinetochore locations (third row), the tracked spindle with bounding boxes illustrates the spindle tracking outcome, (fourth row) the tracked GFP with bounding boxes (red, marked by yellow arrow) depicts the kinetochores being tracked, with bounding boxes delineating the track location (fifth row), and the tracking mask for the GFP signals showcase kinetochore segmentation restricted to the mitotic cell (sixth row). (B) Magnified images highlighted with yellow arrows in Panel A showcase sister kinetochores with two particle IDs (1 and 2) segregated into four kinetochores (1,2,3 and 4). (C) Bar chart showing the accuracy of kinetochore tracking for chromosome 16 evaluated by five assessors, as indicated, for four different movies as those shown in panel A. (D) Bar chart highlighting the combined tracking performance for movies of cells with kinetochore marker for chromosome 16 and chromosome 1, offering insights into the efficacy and reliability of the tracking methodology across different chromosome markers. (E) The table shows sample sizes in panels C, D and Supplementary Fig A2.A. Movie ID description relates the number of tracked kinetochore particles (N), time frames (T) and total number of consecutive events assessed for determining kinetochore tracking accuracies (k). Scale bar: 10 μm.

A watershed segmentation method with object area thresholding is first applied before tracking kinetochores using a Hungarian algorithm [15]. Kinetochores were numbered (Object 1 to 4, referring to two pairs of sister kinetochores) and tracked through anaphase when the two kinetochore pairs underwent segregation into four kinetochores.

Kinetochore object identification and tracking were assessed as correct or incorrect object numbering through consecutive images. Two naive and three experienced users assessed whether object numbers were continuously tracked in four separate movies. Accuracy of tracking shows that in 3 of 4 movies that presented clear signals, kinetochores were correctly tracked through prometaphase, metaphase and anaphase stages (Fig 3.B).

To implement the kinetochore tracker in a more challenging environment, we tested its ability to track an increased number of particles by choosing a second CRISPR guide RNA probe that detected 8 objects instead of 4 by detecting 2 sets of chromosome pairs (Fig A1.A). In these datasets, we observed one set of centromere signals is brighter than the other making the watershed segmentation somewhat difficult (Fig A1.B).

We found an accuracy of 80 to 90 percent in the tracking of correct sister kinetochore particles despite the complexity of these data sets with lower-intensity artefacts (see blue arrow for the challenging low-intensity artefact, Fig A1). Thus intensity-based and spindle mask-constrained watershed segmentation worked satisfactorily to identify and track kinetochores in two different settings.

Combining the accuracies detecting kinetochore particles corresponding to two different chromosome markers in both datasets across over 153 frames (Fig 3.C) shows that 90.83% of kinetochore objects are correctly tracked in the images. This comparative analysis across different chromosomes underscores the tool’s versatility and potential application in broader genetic studies. Thus manual evaluation of computational outcomes shows that the kinetochore tracker works robustly before and after chromosome segregation.

### 2.4 Evaluating the integration of Multi-SpinX output as the input for SpinX

SpinX software has both spindle and cortex segmentation modules. Multi-SpinX was developed to support time-lapse movies of a cell population by isolating single spindles and corresponding brightfield cropped images of the cell cortex. We tested the extent to which SpinX could correctly process the cropped images of spindles and cortex (output of Multi-SpinX). This was important since SpinX was originally trained on metaphase spindles, while Multi-SpinX introduced here can identify prometaphase to late telophase spindles. As expected, late telophase cells and associated spindle fibres although isolated by Multi-SpinX were not detected by the spindle or cortex segmentation modules of SpinX. However, prometaphase and metaphase spindles and cortex were both detected normally in H1299* cells labelled with EB3-mKate2 marker (Fig 4.A). Similarly, during metaphase to anaphase transition in RPE1 cells labelled with SiR-Tubulin dye, the expanding anaphase spindle was selectively segmented by SpinX through time in all consecutive frames; no interphase microtubule fibres were picked by the AI-guided tool (Purple arrow, Fig 4.B).

**Fig. 4:**
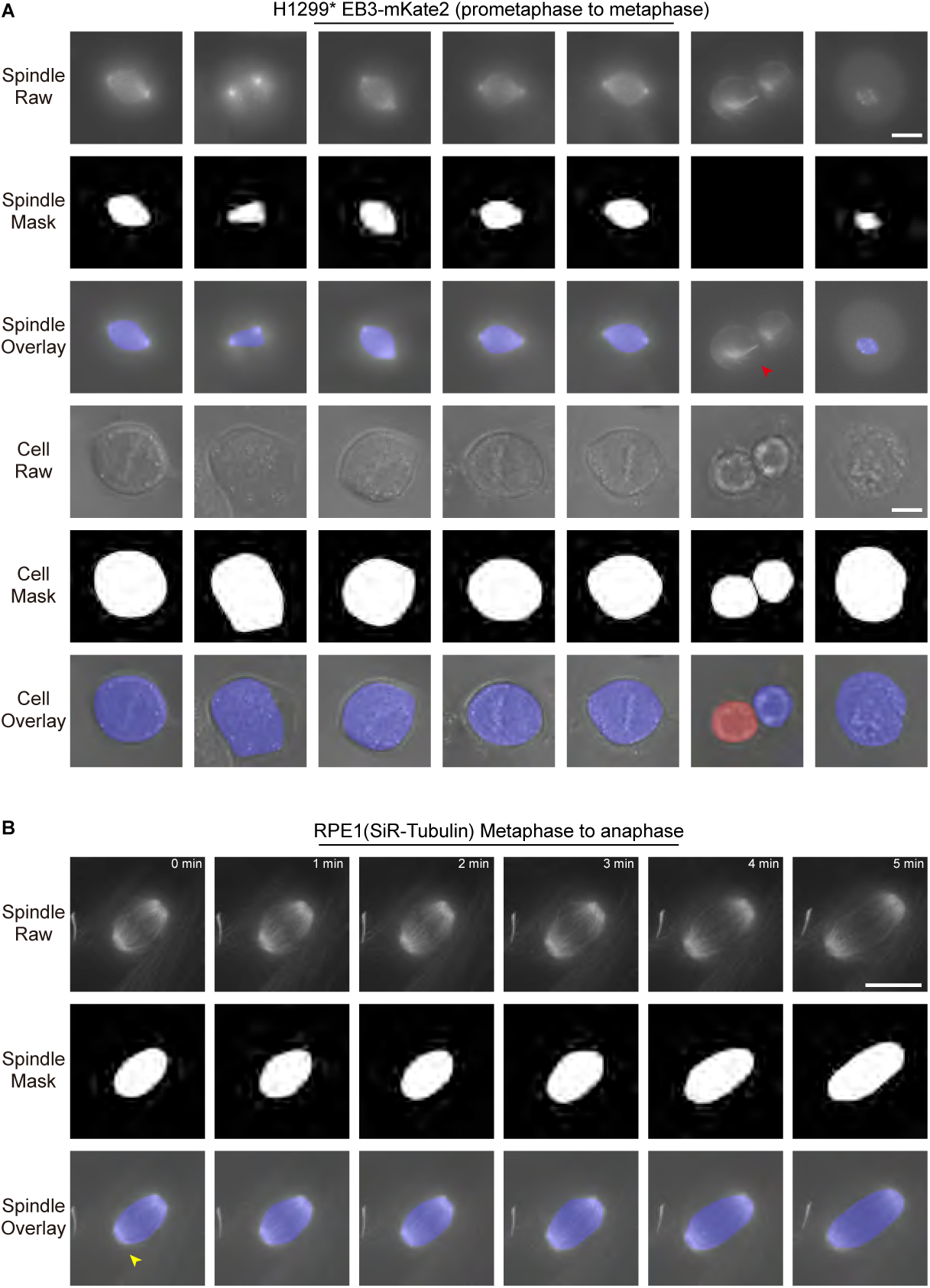
Integration of Multi-SpinX with SpinX’s AI guided models of spindle and cell cortex (A) Sample cropped images for H1299^*^ EB3-mKate2 cell line and their SpinX segmentation inference output for movie 5 frame 1. The original images, their SpinX segmentation mask and the overlay visualisation are displayed in the first, second and third rows, respectively. The original images, SpinX segmentation mask and the overlay visualisation for their corresponding cell cortex are displayed in the fourth, fifth and sixth rows, respectively. The cytokinetic cell (red arrow) was correctly excluded by SpinX segmentation tool as expected. (B) Representative cropped time-lapse images for RPE-1 cells treated with SiR-Tubulin show metaphase to anaphase transition. The original spindle images in consecutive frames are shown in the first row, their SpinX segmentation mask and overlay result are listed respectively in the second and third rows. Interphase microtubule (yellow arrow) excluded by SpinX segmentation tool as expected.

Taking advantage of the successful identification of anaphase spindles by Multi-SpinX, we now tested whether anaphase spindles can be segmented by SpinX. This is important because the AI-guided SpinX software uses a Deep Learning model that was originally trained on metaphase spindles [7], so we assessed the extent to which anaphase A and anaphase B spindles can be accurately segmented by SpinX. Both anaphase A and B spindles in high-resolution images of the RPE1 SiR-Tubulin dataset (60x magnification and images acquired once every minute) could be tracked accurately (Fig 4.B). However, anaphase spindles in low-resolution images of the H1299 dataset (40x magnification) were only occasionally reliable for segmentation via SpinX (Fig A2.B). These findings show that although SpinX segments both low and high-resolution images of metaphase spindles very well [7], this is not the case for anaphase spindles. On the other hand, Multi-SpinX identified both metaphase and anaphase spindles reliably (Fig 2.D), highlighting the complementary strengths of conventional and AI-guided object-tracking tools.

As expected, the correctly segmented metaphase spindle and cortex images could be 3D-modelled for spindle pole positions and spindle centroid within the mitotic cell of H1299 cells (Fig 5.A). In addition to supporting the analysis of multiple spindles encountered in multi-cellular environments, the Multi-SpinX framework can also support the tracking of spindles through different stages of mitosis, making SpinX a versatile tracking software for dividing cells.

**Fig. 5:**
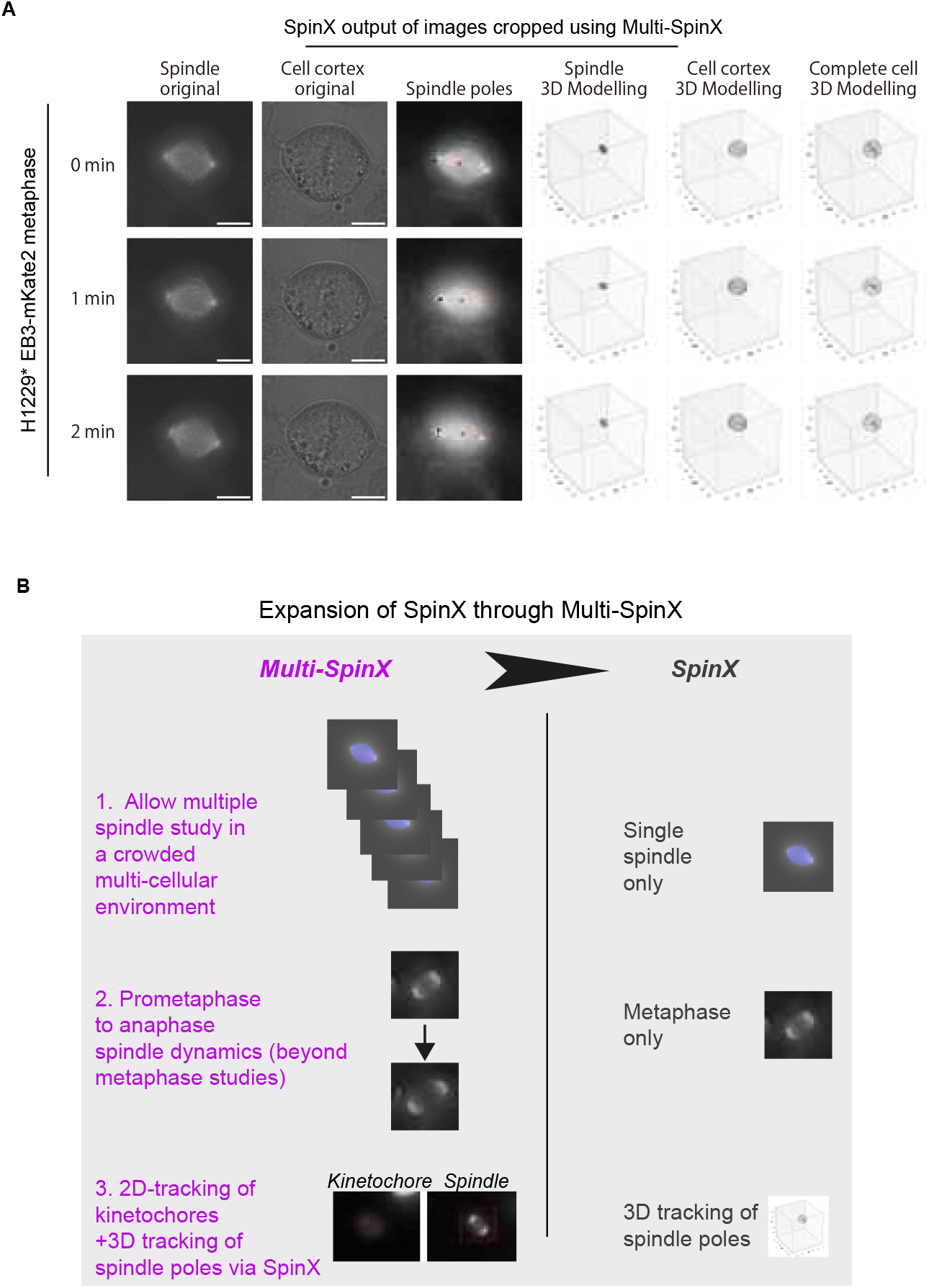
The Multi-SpinX framework makes SpinX a versatile tool for tracking multiple spindles and objects within spindles. (A) Examples of SpinX modelled 3D spindle movements using time-lapse movies of H1299^*^ EB3-mKate2 cells. The crops show one Z-slice of a single spindle and corresponding cortex image Z-stack, from three consecutive frames, employed as the input of the SpinX 3D-spindle modelling module. Spindle pole positions (red and black) and centroids (orange) pseudocoloured via SpinX. (B) Illustration showing how Multi-SpinX strengthens SpinX’s ability to track and model spindle movements in 3D. Multi-SpinX adds at least three new features to SpinX: track spindles in a multi-cellular environment, segment beyond metaphase (prometaphase to anaphase spindles) and track kinetochores within spindle objects.

## 3 Discussion

Here we develop and assess an automated image analysis framework, Multi-SpinX, to significantly extend the applications of the AI-guided spindle tracking software, SpinX. This new extension of SpinX enables 3D-tracking of spindles in time-lapse movies of multi-cellular environments, beyond metaphase into anaphase, and 2D-tracking of kinetochores (new objects) within the larger spindle structure (Fig 5.B). We show the strengths of human-in-the-loop during the process of optimisation and assessment of tracking outcomes, in addition to making SpinX a versatile tool for cell division studies through the Multi-SpinX framework.

The advantage of watershed segmentation for identifying objects like the mitotic spindle is that it can help detect spindles of multiple shapes as they evolve from early to late mitosis (prometaphase to late anaphase). Although our AI-guided SpinX software was trained on metaphase spindles, here we report SpinX’s capacity to segment anaphase spindles in RPE1 cells and demonstrate that the integrated framework can help assess both metaphase and anaphase spindles robustly. Additionally, being able to track metaphase to anaphase to cytokinesis transitions through bounding boxes can significantly reduce the time spent analysing mitotic manually and facilitate the automated linking of mitotic phases with spindle positions which can accelerate future studies of spindle orientation mechanisms [20–22].

Unlike the Kalman Filter and Particle Filter for tracking used in navigation and computer vision systems, extended to dense biological systems [23, 24], which weighs particles on a likelihood score and then propagates them through a motion model to predict their future position, here we successfully implement a Hungarian algorithm that simply takes advantage of the spindle’s and kinetochore’s limited motion since these objects are trapped within a cell cortex. Although the tracking of objects within Multi-SpinX, like other tracking pipelines [5, 8], occurs in 2D, SpinX can model and track individual spindles in 3D, bringing out the strengths of both conventional and AI-guided image processing pipelines.

Like the spindle tracker, the kinetochore tracker can correctly assign object numbers (indicative of correct tracking) through the process of chromosome segregation from early prometaphase to anaphase. Although kinetochore trackers to track metaphase-anaphase transition exist [25, 26], here we track kinetochores labelled by specific centromeric labels which reduces the complexity of crowded particle tracks and allows the successful probing of early prometaphase kinetochore movements and position within the spindle. Thus, the kinetochore tracker along with SpinX can be a powerful tool to assess the rate of chromosome oscillations and segregation in relation to spindle pole movement in 3D, facilitating careful studies of mechanisms that influence both spindle pole and kinetochore function [27–29]. Together, the multi-spindle and kinetochore trackers of Multi-SpinX provide an advanced framework for automated tracking of spindle and chromosome dynamics and their fates in multicellular environments.

## 4 Materials and methods

### 4.1 Data acquisition

#### 4.1.1 H1299^*^ EB3-mKate2 (H1299 *π*-EB1-EGFP/EB3-mKate2) cell line

##### Cell line and culture conditions

The establishment of H1299 *π*-EB1-EGFP (EGFP-Zdk1-EB1C)/EB3-mKate2 cell line, referred to here as H1299^*^ EB3-mKate2, to track spindles, will be discussed elsewhere. H1299^*^ mKate2 cells are cultured in Gibco™ RPMI 1640 medium (FisherScientific™, 11530586) supplemented with Fetal Bovine Serum (FBS; ThermoFisher™, 10270106), 1% penicillin-streptomycin (ThermoFisher, 15140122), 1% MEM Non-Essential Amino Acids Solution (ThermoFisher™, 11140050) and 0.1% Fungizone (ThermoFisher™, 11510496). Cell lines were cultured as a monolayer at 37°C and 5% CO_2_.

##### Live-cell imaging

For live-imaging of H1299^*^ EB3-mKate2, cells were grown in 8-well coverslip dishes (Thistle Scientific, 80806). During imaging, cells were incubated in Leibovitz’s L15 medium (ThermoFisher, 11415064). SiR-DNA dye [30] (Spirochrome, SC007; 1 *μ*M), DMSO (Sigma-Aldrich, D8418) and Ciliobrevin D (CilioDi 5 *μ*M) (Sigma-Aldrich, 250401) were added just before imaging. Live imaging was carried out in a chamber at 37°C using a Deltavision Core (Applied Precision™) microscope equipped with a CoolSnap HQ2 (Photometrics) camera. An Olympus 40x/NA0.95 UPlanSApo air/dry objective was used along with a Quad DAPI/GFP/TRITC/Cy5 polychronic. Movies consisted of 5 z-slices (2 *μ*m spacing) with 5-minute intervals reaching a total time of 4 hrs. Time-lapse images of cells treated with DMSO or CilioDi were pooled into a single dataset.

#### 4.1.2 RPE-1 cell line

##### Cell line and culture conditions

The establishment of the RPE-1 cell line, in which chromosome 1 or 16 is labelled with GFP, will be described elsewhere. Briefly, pericentromeric repeat sequences on each chromosome [31] were visualised by the CRISPR labelling system using nuclease-inactive Cas9 (dCas9) [32]. Cells were grown at 37 °C in a 5% CO2 atmosphere in DMEM (Nacalai), supplemented with 10% fetal bovine serum.

##### Live-cell imaging

Cells were grown in glass chambers (Thermo Fisher Scientific). Five hours before imaging, the medium was changed to prewarmed Leibovitz’s L-15 medium (Thermo Fisher Scientific) supplemented with 20% FBS and 20 mM HEPES, pH 7.0, containing SPY555-DNA (1:10,000, Spirochrome, SC201) and 500 nM SiR-tubulin (Spirochrome, SC002) to visualize chromosomes and microtubules, respectively. The recording was performed in a temperature-controlled incubator at 37 °C. Z-series of eleven sections in 1 *μ*m increments were acquired every 1 min. Images were obtained using an IX-71 inverted microscope (Olympus) controlled by DeltaVision softWoRx (GE Healthcare) using a 60x 1.42 NA UPlanS Apochromat objective lens (Olympus). Images were subjected to deconvolution.

### 4.2 Multi-SpinX pipelines

The Multi-SpinX tracker is designed to monitor the dynamics of multiple spindles in a population of cells across sequential time frames. The first step of the multi-spindle tracker is preprocessing, where maximum projection is first applied to the multi-stacked movie before a min-max normalisation for contrast adjustment. The tracker then employs watershed segmentation on the preprocessed movie that identifies the presence of multiple spindles within each time frame. Following detection, a Hungarian algorithm with Euclidean distance as the loss is used for object matching across consecutive time frames (see Table B1 for hyperparameter setting). All detected spindles are isolated with precision using bounding boxes. Bounding boxes are applied to the cell cortex channel before the generation of cropped spindle and cell cortex image pairs for all the detected spindles across time. The cropped spindle and cell cortex image pairs then are run through SpinX [7] for 3D spindle modelling. This step takes z-slice information and reconstructs the two-dimensional images into detailed three-dimensional representations of each spindle, providing a richer and more comprehensive view of spindle morphology and dynamics [7].

Complementing the multi-spindle analysis, the kinetochore tracker embarks on a focused mission to detect and track kinetochores within the spindles of interest. Spindles are first detected with the multi-spindle tracker, a watershed segmentation is then used for the kinetochore segmentation on the GFP channel on top of spindle detection(see Table B2 for hyperparameter setting). Following the segmentation, kinetochores are tracked across successive time frames using the Hungarian algorithm to ensure kinetochores are accurately paired from one frame to the next despite their dynamic movements. This tracking capability is vital, as it would allow paired monitoring of kinetochore/centromere behaviour throughout mitosis, during chromosome capture and segregation.

### 4.3 Benchmarking and analysis

Original live-cell movies of H1299^*^ EB3-mKate2 and processed movies with bounding boxes were analysed by expert and naive users. Three naïve and expert users analysed spindle tracking data, while GFP-labelled kinetochore tracking data was analysed by five naïve and expert users. All movies were viewed and manually assessed using Fiji [33] The assessments of spindle and kinetochore tracking were performed using EB3-mKate2 or GFP-labelled centromere signals, respectively. The maximum intensity of Z-slices was determined using the Z-project function. For the spindle tracking data, the assessment was performed in two iterations: the first iteration was primarily to identify tracking-related issues (details in next section), which were addressed before the second iteration where the accuracy of the multi-SpinX tracker was manually assessed. For the kinetochore tracking data, in RPE1 cell lines with either chromosome 1 or 16 GFP-tagged, the assessment was performed in a single iteration.

#### Multi-SpinX assessment

In the first iteration of the assessment, users visually categorised spindles as correctly or incorrectly tracked through consecutive time frames with corresponding colour codes (eg., low-intensity spindles, spindles close to the image boundary, and spindles beyond telophase). The second iteration of the assessment categorised spindles with a unique bounding box ID to facilitate accurate tracking assessments.

#### Metaphase and anaphase spindle tracking analysis

To analyse metaphase and anaphase spindle tracking, three naïve and expert assessors manually counted the number of metaphase or anaphase spindles missing bounding boxes and thus not being tracked. Average percentage values of all 3 assessors were calculated. Only correct bipolar spindles were considered. Multi-polar or overlapping spindles were exempt from analysis.

#### Kinetochore tracking analysis

To assess kinetochore tracking during chromosome segregation, the splitting of two bounding boxes (corresponding to two sister kinetochore pairs) into four bounding boxes (corresponding to segregated four sister chromatids) were assessed as ‘incorrectly’ tracked or ‘artefacts’ suggesting a mis-tracked signal. For each time frame of the movie, numbers of correctly and incorrectly tracked kinetochore pairs or kinetochores were counted and the average percentage was calculated across values from all five assessors.

### 4.4 Statistical analysis

Data from each assessor was compiled and an average percentage of correctly and incorrectly tracked objects (metaphase and anaphase spindles, and kinetochores), in each movie was determined. Statistical tests were performed in Graph Pad Prism using two-tailed, Paired T-tests as well as multiple paired T-tests across movies. The normality of data was confirmed using Q-Q plot. Statistical tests were performed on a significance level of P*≤*0.01 or P*≤*0.05. For P values, the following convention holds: not significant (n.s.) with P*>*0.05, (*\**) with P*≤*0.05, (*\*\**) with P*≤*0.01, (*\*\*\**) with P*≤*0.001 and (*\*\*\*\**) with P*≤*0.0001.

## Data availability

Raw data are available on Zenodo; Image analysis codes and manual assessment outcomes are all available in Github: https://github.com/Draviam-lab

## Author contributions

V.M.D and B.C. drafted the manuscript together. B.C, V.M.D. and C.E. edited sections using comments from M.C. B.C. contributed to Fig 1 with support from V.M.D.; B.C., C.E, M.C, S.J. and A.M. contributed to Fig 2; B.C., K.T., K.K., S.J. and M.C, contributed to Fig 3 and Fig A1 and Fig A2.A; B.C contributed to Figs 4. B.C and V.M.D contributed to 5. C.E., analysed images in Fig A2.B. K.K., K.T., B.C. and V.M.D contributed to Fig A1.B.

## Acknowledgments

We thank Dr Torsten Wittman’s Lab for sharing their H1299 EGFP-Zdk1-EB1C (*π*-EB1) parent cell line. We thank members of the group of V.M.D. for useful discussions. We acknowledge funding support from the Biotechnology and Biological Sciences Research Council (BBSRC) and InnovateUK to V.M.D. (BB/R01003X/1, BB/T017716/1, BB/W002698/1, and BB/X511067/1 and KTP012502), B.C. (BB/X511067/1 and InnovateUK KTP012502), and C.E. (LIDo-iCASE studentship BB/T008709/1). A joint studentship from QMUL and Zeiss funds M.C. We acknowledge preliminary tracking efforts by Mohammad Javad Shojaei (supported by InnovateUK KTP012502 funding to VMD) which is not considered because of its incomplete and inconclusive nature. B.C. was a Knowledge Transfer Partnership associate collaborating with ZEISS UK, supported by BBSRC FTMA (2023) and Regional England Investment Fund (2024).

## Appendix A Supplementary figures

**Fig. A1:**
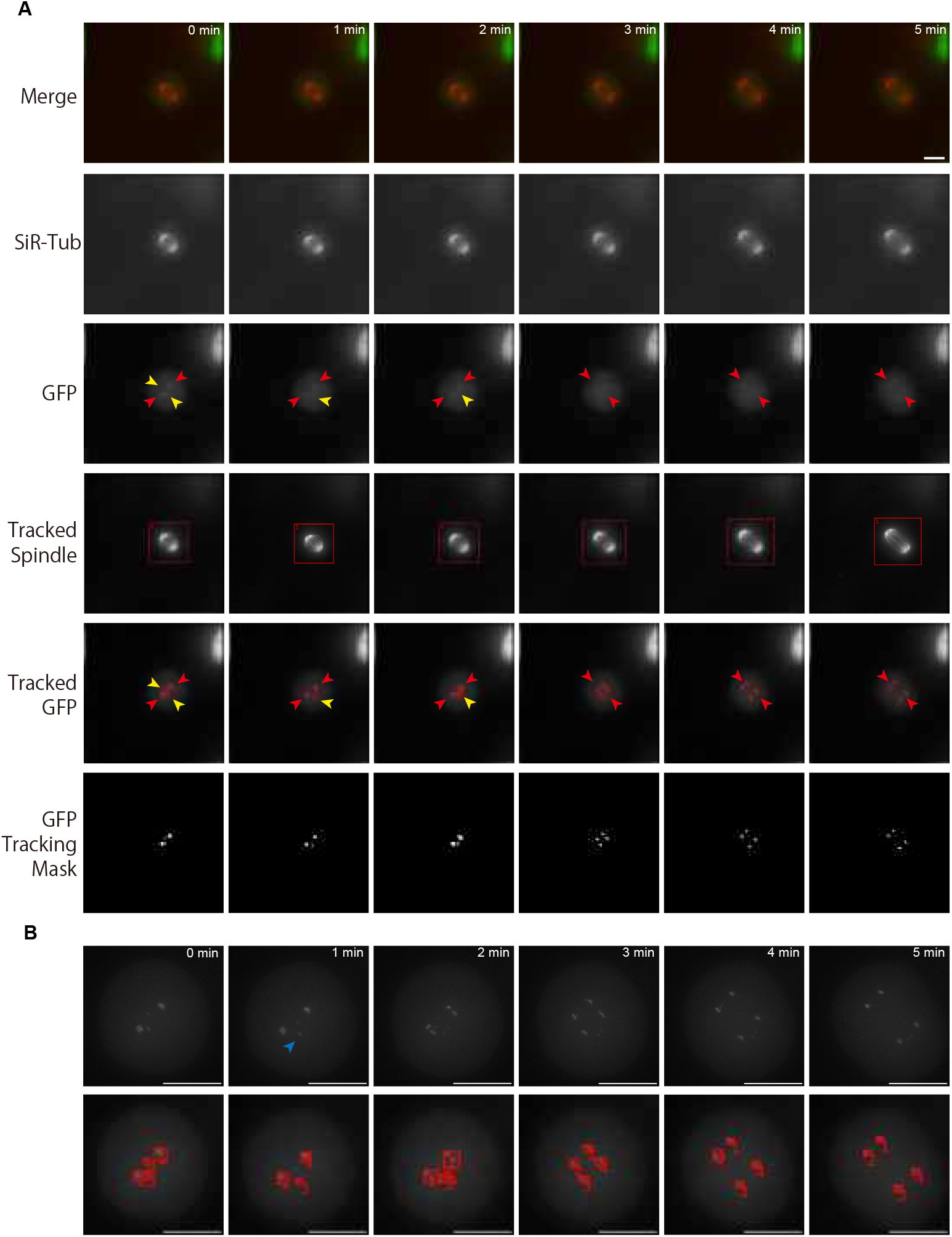
Sample images for the CRISRP engineered kinetochore tracker labelling chromosome 1 in RPE1 cells. (A) Time-lapse images of consecutive frames illustrate the complex process of kinetochore tracking overlaid with spindle tracking. The frames are organised as follows: the original movie as a merge of the spindle (red) and kinetochore marker (centromere marker for chromosome 1, in green) displays the representative consecutive frames (first row), SiR-Tubulin (SiR-Tub) labelled channel highlights the spindle structure (second row), GFP channel highlights the kinetochore locations (third row, red, yellow and red arrows marked), the tracked spindle with bounding boxes illustrates the spindle tracking outcome, (fourth row) the tracked GFP with bounding boxes (red, yellow and red arrows marked) depicts the kinetochores being tracked, with bounding boxes delineating the track location (fifth row), and the tracking mask for the GFP signals showcase kinetochore segmentation restricted to the mitotic cell (sixth row). (B) This panel shows the magnified images highlighted with red and yellow arrows in Panel A. Red arrows showcase the dominant sister kinetochores with two particle IDs segregated into four kinetochores for chromosome 1. Yellow arrows show another set of centromere signals which is weaker compared to the dominant signals, but these are brighter than in the chromosome 16-labelled movies making the watershed segmentation somewhat difficult. Scale bar: 10 μm.

**Fig. A2:**
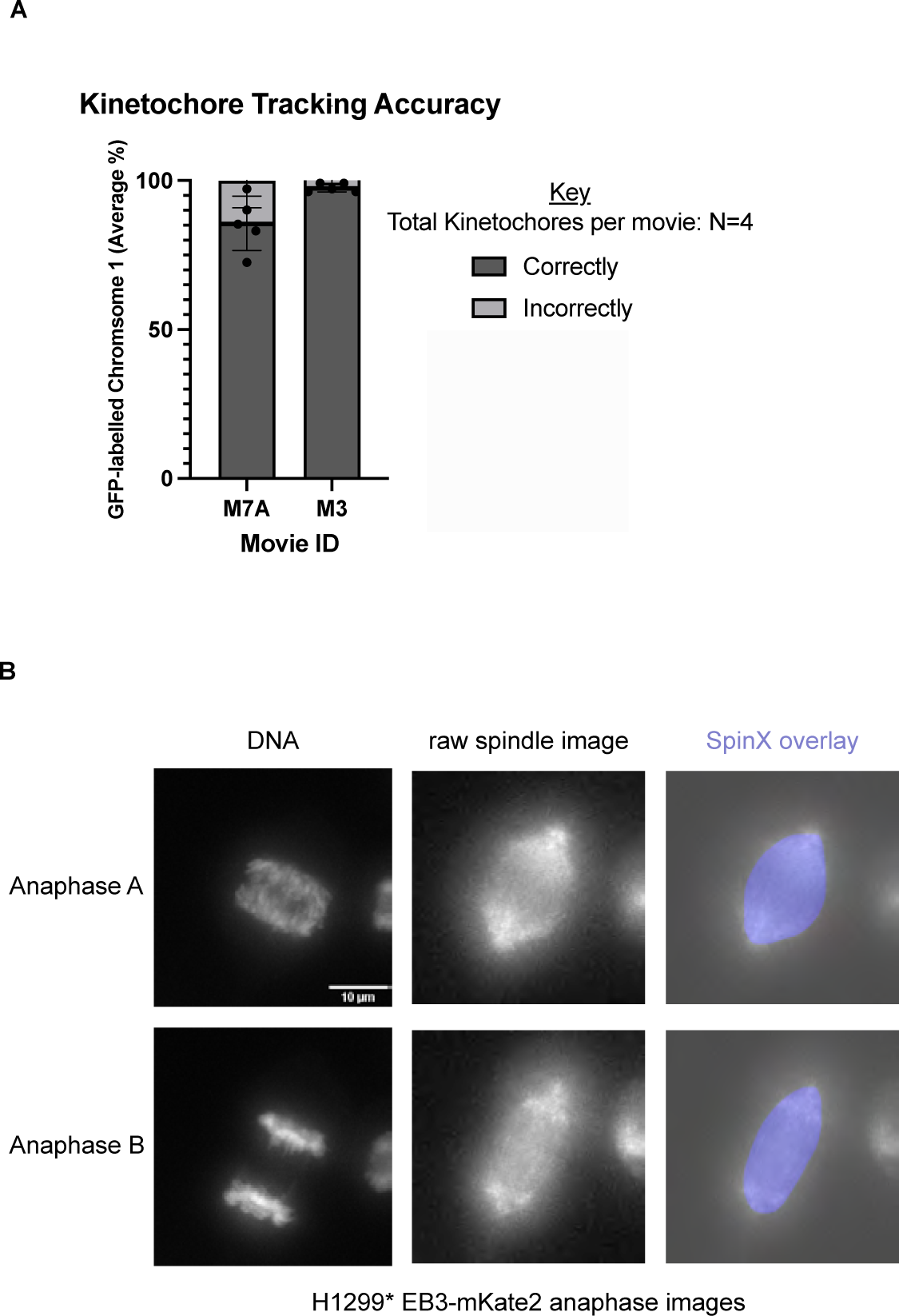
Two complex image datasets to test the boundaries of Multi-SpinX based tracking of smaller particles within the larger spindle (kinetochore) and low spatial and temporal resolution images of anaphase spindles (A) Bar chart showing the accuracy of kinetochore tracking for chromosome 1 in RPE1 cells which presented a new complexity of additional artefactual low-intensity particles (Data related to Fig A1). Two movies were evaluated by five assessors, Sample sizes of movie frames are indicated in Fig 4.E. (B) Representative Multi-SpinX crops of low-resolution spindle images of the H1299* EB3-mKate2 cell line (middle row) show successful identification of Anaphase spindles (left row showing chromosome segregation status), and SpinX implementation outcome (overlay, right column). Very few anaphase images could be spotted in this dataset of movies taken once every 5 minutes, analysis related to Fig 2. Scale bar: 10 μm.

## Appendix B Supplementary tables

**Table B1:** Hyperparameter settings for the multi-spindle tracker.

| ID | Frames | Watershed lower and higher marker values | Auto adjust enabled |
| --- | --- | --- | --- |
| 01 | 1-25 | 0.30, 0.40 | n |
| 04 | 1-9 | 0.30, 0.40 | n |
| 05 | 1-11 (05a), 34-49 (05b) | 0.30, 0.40 | y |
| 06 | 1-16 (06a), 27-31 (06b) | 0.30, 0.40 | n |
| 08 | 1-24 (08a), 33-41 (08b) | 0.30, 0.40 | n |
| 09 | 1-49 | 0.30, 0.40 | n |
| 10 | 1-49 | 0.30, 0.40 | n |

**Table B2:**
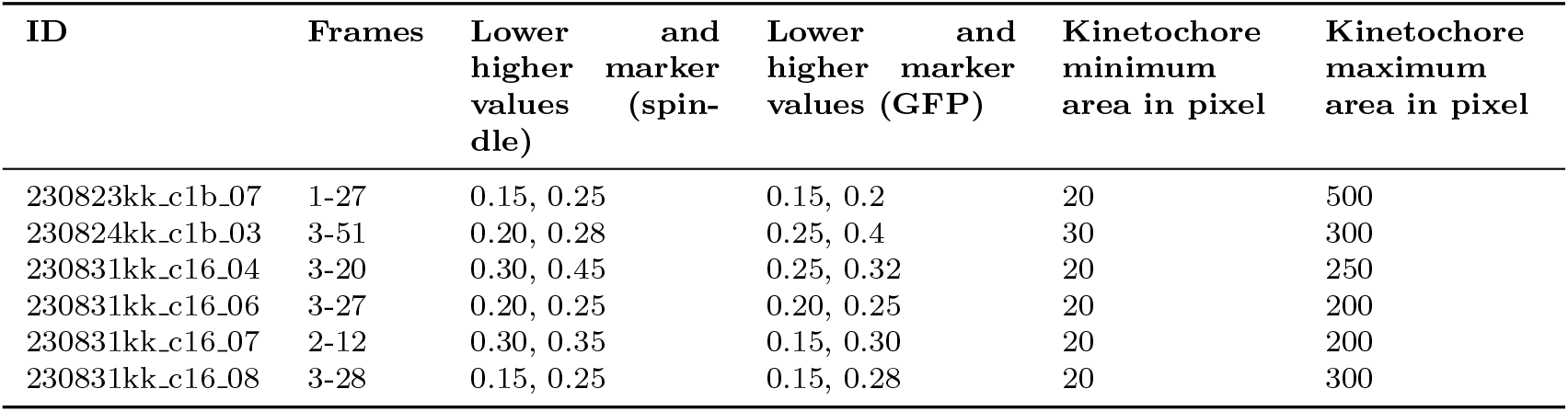
Hyperparameter settings for the kinetochore tracker.

